# Detection of zoonotic *Cryptosporidium parvum* in invasive beavers from southern Tierra del Fuego, Chile

**DOI:** 10.1101/2025.11.13.688090

**Authors:** Paola Larrauri-Aguilar, Gabriela Contreras, Gonzalo Cabrera, Cristóbal Arredondo, Catherine Dougnac, Fernán Federici, Cristóbal Briceño

## Abstract

Since their introduction to Tierra del Fuego in 1946, invasive beavers have expanded their population and caused extensive environmental degradation across the sub-Antarctic region. Beaver dams have profoundly impacted peatlands, mature sub-Antarctic forests, and watercourses. Here, we report a pathogen in invasive beavers in Patagonia and discuss their potential role as a reservoir of waterborne zoonotic pathogens. We detected *Cryptosporidium* through microscopy using the Ziehl-Neelsen staining technique in beaver collected from southern Tierra del Fuego. Out of the 105 beavers examined, three tested positive to optical microscopy, which were then identified molecularly as *Cryptosporidium parvum* by sequence analysis of the 18S ribosomal DNA (rDNA). Given the potential impacts of *Cryptosporidium parvum* upon native species, wildlife, and human health, this finding highlights the potential risks associated with the ongoing expansion of beavers into other areas of the sub-Antarctic region. A potential role of invasive beavers as reservoirs of zoonotic *Cryptosporidium* spp. in Patagonia emphasizes the unknown consequences of human-introduced exotic species on pristine ecosystems.

## Introduction

*Cryptosporidium* spp. are waterborne pathogens that cause gastrointestinal disease [1]. *Cryptosporidium* infects several animal groups, including cattle, poultry, rodents, domestic pets, and humans [2]. The infective stage is a thick-walled oocyst, 4-6 μm in diameter, and ingestion of as few as 1 to 10 oocysts can be sufficient to cause infection [3], [4]. The trilaminar oocyst wall enables survival outside the host and, upon ingestion, releases sporozoites that infect the epithelial cells lining the host’s gastrointestinal mucosa [5]. This adaptation allows the oocyst to remain viable in the environment for up to 16 months [1]. Oocysts can endure ultraviolet (UV) light and chlorine in high parasitic loads [3], but not alkalinity, desiccation and ozone exposure [6], [7].

Oocysts are excreted in feces and transmitted via the fecal-oral route. This transmission occurs through direct contact with infected hosts or indirectly through the consumption of contaminated water and food [2]. Sewage and watercourses in livestock-dense areas show higher oocyst levels than regions with minimal human intervention [6]. Oocyst contamination of water supplies is driven by flow, seasonal precipitation and runoffs [6], [8], [9] as they mobilize oocysts from fecal sources and transport them to watersheds. Another key factor is domestic animals, wildlife and invasive species acting as pathogen reservoirs and interacting with these environments [10], [11].

*Cryptosporidium* spp. accounts for most waterborne protozoan outbreaks worldwide between 1984 and 2003 (165 of 325, 50.8%), with 50.3% of cryptosporidiosis cases associated with contaminated recreational water [12]An updated review reported that 63% (239/381) of worldwide outbreaks between 2004 and 2014 were attributed to *Cryptosporidium* spp. [13]. *Cryptosporidium parvum* is strongly associated with mortality in children and immunocompromised patients and causes diarrhea in adults [1]. In Latin America, *C. parvum* shows the widest geographical distribution and circulation, largely attributed to primary zoonotic sources such as cattle and sheep [14]. However, wildlife reservoirs remain understudied in southern ecosystems.

Invasive alien species (IAS) contribute to biodiversity loss [15] and species extinction [16]. They cause biotic homogenization and ecosystem transformation, including isolated environments like archipelagos, which are more vulnerable to their impact [17]. Such changes may facilitate disease onset by establishing novel links between emerging pathogens and wildlife hosts [10], [18]. As an IAS expands, pathogen-host interactions may increase the risk of zoonotic spillovers, influenced by environmental factors, epidemiology, host susceptibility, and pathogen ecology [6], [15].

Tierra del Fuego (TDF) archipelago, located at the southernmost tip of Chile and Argentina, is a remote ecosystem with high endemicity. Once considered one of the world’s most pristine areas, this region was drastically altered in the second half of the 19th century due to extensive sheep farming [19], [20]. Today, it faces threats from extensive cattle farming and the oil industry [21]. Furthermore, the archipelago’s unique and diverse ecosystems are continually threatened by IAS, which represent two-thirds of the archipelago’s species richness. This has the potential to disrupt the region’s ecological balance [17], [20].

Among IAS, the North American Beaver (*Castor canadensis*) has a significant impact on peatlands, mature sub-Antarctic forests, and watercourses [20]. In 1946, ten reproductive pairs from Canada were introduced to the Argentinian side of the TDF Island. This introduction expanded its distribution onto the continent and northward, becoming a population of estimated 100,000 individuals [22]. The abundance of resources and the absence of natural predators and competitors facilitated an expansion rate of 2-6 linear km/year [23], [24]. Moreover, beaver colonies in TDF are reportedly higher in density (4 to 5 colonies per km^2^) than those reported in North America (up to 3 colonies/km^2^) [25].

Beavers are semi-aquatic animals that exhibit robust social structure; they inhabit watersheds and build dams at dawn and dusk [26]. During the first 2-3 days post-birth, mothers ingest their kits’ feces. Subsequently, kits defecate into the water around the burrow [26], contributing to sediment accumulation and watershed pollution throughout their lives. Beavers have been reported as reservoirs of *Giardia* (assemblage A) and *Cryptosporidium* (*C. ubiquitum* and *Cryptosporidium* sp. beaver genotype) in North America using immunofluorescence microscopy (IFA), PCR analysis and sequencing [27], [28], which would suggest their potential contribution in the dissemination of waterborne pathogens in Patagonia [9].

Beaver activity also creates habitats suitable for IAS, including exotic plants, salmonids, and semi- aquatic mammals like muskrats (*Ondatra zibethicus*), and American minks (*Neogale vison*), the latter being an opportunistic carnivore [20]. Interactions between these sympatric species in North America and Patagonia produce an invasive meltdown favoring synergistic dynamics [29], which may increase pathogen transmission due to their close interactions.

We hypothesize that North American beavers, introduced to TDF eight decades ago, harbor *Cryptosporidium* spp. and represent an undocumented reservoir for waterborne pathogens in this region. To elucidate environmental contamination pathways and watershed pathogen dynamics within a One-Health framework, we (i) estimate *Cryptosporidium* sp. prevalence in beaver populations, and (ii) characterize oocysts morphology and species using microscopy and DNA sequencing.

***Graphical abstract***. The influence of beaver activity on animal and environmental health in Tierra del Fuego. Beavers expose native species (1-3), livestock, domestic animals, and humans (4-6), and other invasive species (7-8) to contaminated water with exudates, potentially facilitating *Cryptosporidium* oocysts dissemination throughout the region. **(1)**: Traro (*Caracara plancus*); **(2)**: Culpeo fox (*Lycalopex culpaeus lycoides*); **(3)**: Guanaco (*Lama guanicoe*); **(4)**: livestock; **(5)**: domestic animals; **(6)**: humans; **(7)**: Muskrat (*Ondatra zibethicus*); **(8)**: American Mink (*Neogale vison*). Created in https://BioRender.com

## Materials and methods

The Global Environmental Facility (GEF) program, involving FAO (UN Food and Agriculture Organization), SAG (Fish and Wildlife Service), CONAF (Protected Natural Park Service), and the Wildlife Conservation Society (WCS) Chile, coordinated a control, prevention, and management project of the beaver population in the 300,000-hectare Karukinka Park, a natural laboratory in TDF, managed by WCS Chile [30]. 110 beavers were captured from three locations on TDF Isla Grande (54°S 69°W), Chile in spring and summer between 2017 and 2019 [31] (**Figure 1**). Necropsies were performed, and fecal samples collected from rectum of 105 beavers. Five of them were dismissed due to intestinal and rectal damage, either due to the impact of the trap or to initial decomposition from submersion, as Conibear traps are typically located in underwater channels [32].

**Figure 1.**
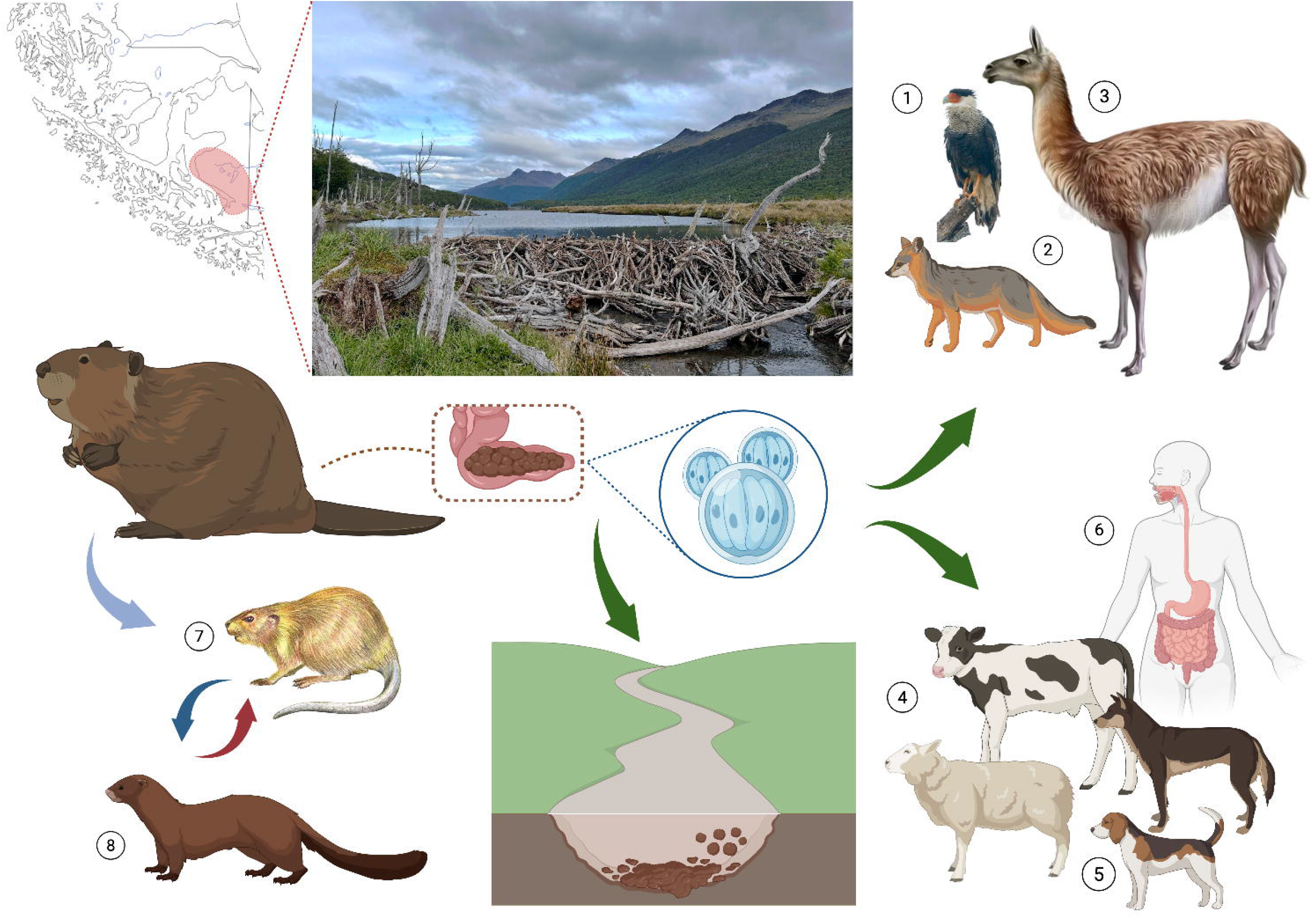
Beaver sampling locations in Tierra del Fuego Archipelago between 2017 and 2019. Beaver sampled locations are depicted with circles, in blue for negatives and red for positive samples of *Cryptosporidium parvum*.

All 105 stool samples (3cm^3^ in volume) were stored in 70% ethanol, transported at room temperature (RT) to Santiago, and kept at 4ºC until processing. Ethanol was decanted by inversion into another sterile tube; each fecal sediment was prepared on one slide and stained using the modified Ziehl-Neelsen method for *Cryptosporidium* oocysts detection through light microscopy (100X). Suspect oocysts showed subspherical morphology, 4 – 6 μm size, fuchsia color, and visible content (four sporozoites) [33].

For DNA extraction, ethanol was decanted, and 200 mg of fecal samples were resuspended in 850 μL Lysis Buffer (NucleoSpin^®^ DNA Stool Mini Kit, Macherey-Nagel). Samples were treated with ten freeze-thaw cycles using liquid nitrogen and a 65ºC water bath for one min each per cycle, vortexed with 0.6–0.8 mm ceramic beads (2mL MN Bead Tube Type A) for 25 min to ensure thick-walled oocysts’ disruption [34]. Genomic DNA was extracted using the NucleoSpin^®^ kit and stored at -20ºC for analysis. Finally, DNA was concentrated 6.6-fold by lyophilization to improve the target detection. A nested PCR protocol targeting the 18S ribosomal RNA (rRNA) gene, a marker distributed across the *Cryptosporidium* genus, was used [34]. This marker features semi- conserved and hypervariable regions used for genus and species identification [35]. This PCR technique enhances detection sensitivity, given low oocyst loads in water and stool [36].

Target sequences were used to build a synthetic template containing regions of the small (SSU/18S) and large (LSU/28S) subunits of the *Cryptosporidium* rRNA gene. Three sequences were selected (Accession Numbers: MZ892388.1, AF108862.1, AF040725.1), keeping only relevant DNA regions for primer compatibility. Sequences were merged using Benchling, synthesized as a gBlock, housed in a pL0R-LacZ cloning vector [37], and cloned into *E. coli* TOP10 cells for supply. A synthetic control was produced Crypto_18S rDNA C+ [38]. Nested PCR was performed using primers described by Briceño [34], that target a well-conserved region within the 18S rRNA gene. 50 μL-volume PCR reactions contained 10 μL of High-Fidelity 5X Buffer, 200 μM each dNTP, 0.4 μM of each primer, 0.5 μL (20 units/ml) of Phusion High- Fidelity DNA Polymerase, 1.66 μL of DNA solution, and molecular-grade water. For the secondary reaction, 1.66 μL of primary product served as template under identical conditions. Thermocycling included initial denaturation at 98ºC for 30 s, followed by 38 cycles consisting of 20 s at 98ºC, 45 s at 65ºC, and 1 min at 72ºC, with a final extension of 7 min at 72ºC for the primary reaction. Secondary reactions used 38 cycles of 20 s at 98ºC, 30 s at 65ºC, and 40 s at 72ºC. PCR products were visualized in 1% agarose gels, using SYBR^®^ SAFE dye (Invitrogen) and an open- source blue LED Mini Transilluminator (IO Rodeo).

552-bp Nested PCR products were inserted into a pL0R-lacZ vector using Type IIS Golden Gate assembly (with SapI and T4 ligase enzymes) for greater stability. Plasmids were cloned using *E. coli* TOP10 chemo-competent cells [37], purified for storage in glycerol (-80ºC), and species were identified by Sanger sequencing using universal primers (UNSp1F and UNSp1R) described by Pollak (2020) [37]. Manually cleaned sequences were aligned and a phylogenetic tree was constructed using the Maximum Likelihood (ML) test and the Tamura-Nei evolutionary method (TN92+G+I) in the MEGA 12 software [39] to compare the PV442491.1 *Cryptosporidium* clone **PL14** 18S rRNA sequence (target) with *Cryptosporidium* spp. sequences listed in **Supplementary Table S1**.

## Results

From 110 trapped beavers, 50% were female, 51% were adults, 20% were kits, and 29% were juveniles. Three beavers tested positive for *Cryptosporidium* spp. by optical microscopy (3/105) using the Ziehl-Neelsen staining technique: two adults (CC52 and 55) and one juvenile (CC53); among these, CC52 and CC53 were females, while CC55 was male. To identify the parasite species, we amplified and sequenced a fragment of the 18S rRNA gene. Our findings confirm zoonotic *C. parvum* in beaver feces.

### Microscopic and molecular detection

Three beaver samples tested positive for *Cryptosporidium* spp. using both microscopy and nested PCR techniques. **Table 1** provides further information of the former three beavers. Among these, both females showed common capture locations (coordinates). **Figure 2** shows the identification of one oocyst per sample through microscopic analysis (CC52 slide shown), and amplicons from nested PCR in agarose gel.

**Table 1.** Characteristics of trapped beavers positive for *Cryptosporidium*.

| **Code** | **Sex** | **Age** | **Weight** | **Location coordinates** | **Condition at capture** |
|---|---|---|---|---|---|
| CC52 | Female | Adult | 19.57 kg | 19 F 0513545 UTM 3975660 (La Paciencia) | Entire animal underwater |
| CC53 | Female | Juvenile | 16.81 kg | 19 F 0513545 UTM 3975660 (La Paciencia) |  |
| CC55 | Male | Adult | 21.61 kg | 19 F 0511726 UTM 3976927 (La Paciencia) |  |

**Figure 2.**
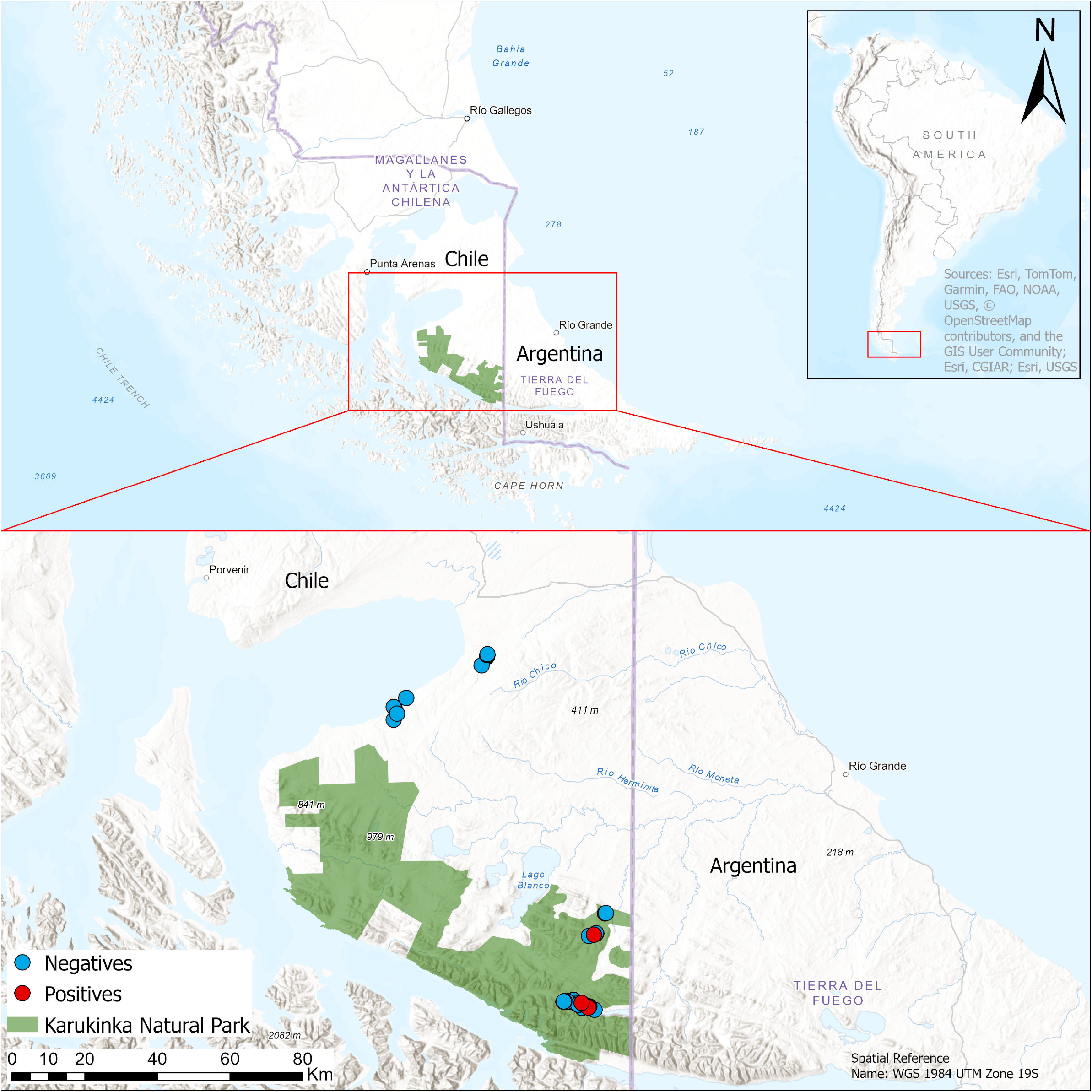
Detection of Cryptosporidium using optical microscopy and nested PCR. (A) (left): Ziehl-Neelsen-stained beaver feces specimen from Tierra del Fuego (spherical fuchsia body, observed with 100X lens). (B) (right): Molecular confirmation of the *Cryptosporidium* 18S rRNA gene (552-bp bands on agarose gel) in three beaver samples (CC52, CC53, CC55; lanes 4, 5, and 6, respectively). Lane 1: pL0R plasmid as negative control (C-); Lane 2: Ladder 100bp (L); Lane 3: synthetic positive control (C+); Lane 7: no template control (NTC).

### Sequence analysis

Alignment of sequenced plasmids yielded a 451-bp 18S rRNA fragment (after trimming primers and uneven chromatograms), available at GenBank as PV442491.1. This sequence showed 100% identity with *C. parvum* verified by phylogenetic analysis using the ML test, TN92 + G + I model and 1,000 bootstrap replicates.

The resulting phylogenetic tree revealed a distinctive *C. parvum* clustering comprising 13 isolates separated from *C. bovis, C. hominis, C. meleagridis* and other species (**Figure 3**).

**Figure 3.**
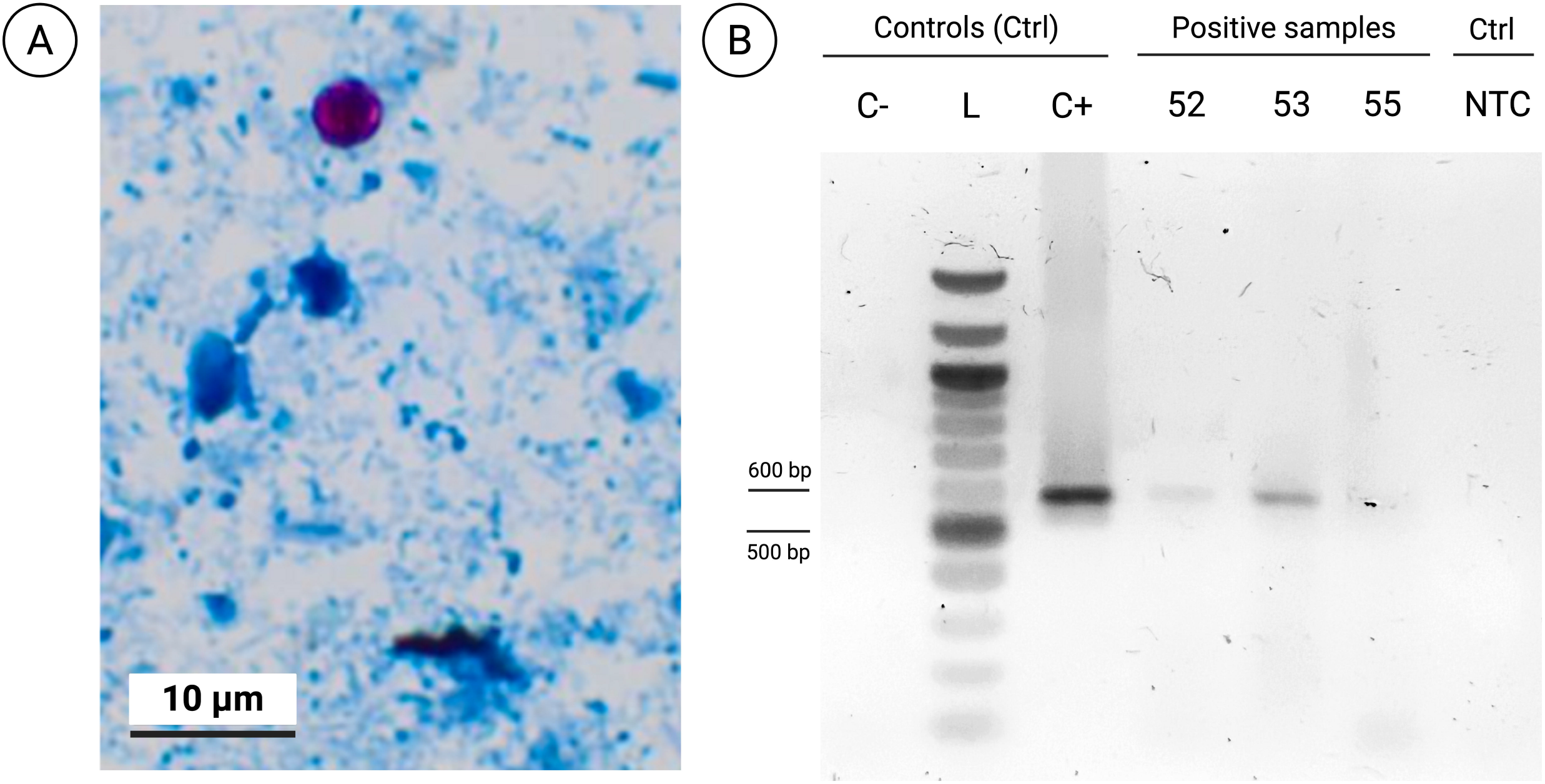
Phylogenetic tree of *C. parvum* strain PV442491.1 based on 18S rRNA sequences. Maximum-likelihood phylogeny constructed using MEGA 12 software with Tamura-Nei (TN92+G+I) model of nucleotide substitution. Sequences aligned with *Clustal W* (alignment length 477 bp, 377 conserved sites, 87 variable sites, 60 parsimony informative sites). GenBank accession numbers and species displayed at terminal nodes. Bootstrap support values from 1000 replicates shown at internal nodes (≥ 70% only). Scale bar indicates 0.5 substitutions per site. The target sequence PV442491.1 (outlined in red) clusters within the *C. parvum* clade, distinctly separated from other sequences. *Plasmodium falciparum* 18S rRNA sequence is used as outgroup.

## Discussion

Our research showed that invasive beavers in TDF carry *C. parvum* oocysts. Although our study cannot establish beavers as definitive *C. parvum* hosts, as immunohistochemical assays on intestinal tissue are still required to confirm an active infection [9], these findings raise urgent questions about their potential role in the dissemination of this parasite. As ecosystem engineers, beavers could potentially contribute to the dissemination of oocysts via sediment and fecal accumulation throughout the basins of the study area and beyond (**Figure 1**). The infection rate observed over two-year sampling period (2017-2019) was relatively low (2.87%), in concordance with studies in North America (USA) reporting a 2.2% to 18.9% prevalence range among American beavers [27].

Our findings showed a very low parasite load in the beaver feces. The conditions and adjustments used in the pre-treatment of samples–thermal shock (freeze-drying) combined with mechanical disruption–and DNA lyophilization as a concentration strategy, were sufficient to detect the parasite DNA through nested PCR targeting the 18S rRNA gene (**Figure 2**). Subsequent phylogenetic analysis confirmed the PV442491.1 sequence as the clinically relevant *C. parvum* (**Figure 3**), a species with low specificity and significant zoonotic potential [12]. Therefore, our findings indicate potential risks for native, exotic and domestic species, as well as humans [9], [40].

To our knowledge, this study provides the first evidence of *Cryptosporidium* presence in invasive beavers in TDF. This parasite had been previously reported in Adélie penguins (*Pygoscelis adeliae*) on Ardly Island, Antarctica (6.59% of 167 samples) [41] and Magellanic penguins (*Spheniscus magellanicus*) [42]. A study from Massachusetts (USA) reported a 3.2% prevalence of *Cryptosporidium* in North American beavers (2/62) using IFA and PCR [43]; both were from kits.

Given that beavers as an IAS alter disease transmission dynamics by creating novel host-parasite assemblages, assessing pathogen occurrence in natural environments is crucial [6], [44]. A context- specific approach to pathogen epidemiology is essential [9]. Surveillance should include waterborne pathogens with a higher prevalence among native species such as guanacos (*Lama guanicoe*), traros (*Caracara plancus*), and Fuegian culpeo foxes (*Lycalopex culpaeus lycoides*) [45] sharing water bodies. Invasive species such as chilla foxes (*Lycalopex griseus*), American minks, and muskrats [29] may also encounter intervening watercourses or vegetation irrigated by beaver dams, or prey on beaver carcasses [23]. Moreover, livestock, domestic animals, and feral dogs (*Canis familiaris*) may spatially overlap with the former (**Graphical abstract**). This dynamism could increase exposure as infected hosts defecate near streams or within dams, thereby acting as reservoirs that support ongoing transmission [27].

These findings call for further research to better understand the role of beavers in *C. parvum* epidemiology. Analyzing a larger, more representative sample of beaver colonies would help elucidate more accurate age-stratified prevalence estimates. Beyond the well-documented impact on sub-Antarctic forests and water systems, their putative role as reservoirs of animal and human pathogens opens a new horizon of threats to both conservation and public health [20]. As beaver populations are migrating northward, coordinated One Health-based surveillance is imperative [22].

## Limitations

### Microscopy vs. molecular characterization of *Cryptosporidium* spp

Microscopy-based detection is the gold standard for detecting *Cryptosporidium*. This method, however, analyzes a single slide per sample, with a smear representing only a minimal fraction of the fecal matter. Given the uneven oocyst distribution in fecal matter and interference from the sample’s physical and organic properties [46], this approach may underestimate the actual parasite load in a sample. In contrast, molecular techniques can confirm findings and identify false negatives, as well as discriminate among species [3]. PCR-based methods are highly recommended as they can detect DNA that remains intact for a long time, ensuring a high detection rate [4]; moreover, nested PCR offers enhanced sensitivity and specificity than conventional PCR. To accurately differentiate parasite loads across samples, it is necessary to incorporate quantitative PCR (qPCR) analysis [9], a method sensitive enough to detect as few as three oocysts [3].

### 18S rRNA vs. gp60 use for molecular characterization of *Cryptosporidium* spp

This study aimed to identify and verify the presence of *Cryptosporidium* spp. in clinical, environmental, and experimental samples. The 18S rRNA gene was used for initial screening and environmental surveillance because of its low interspecific variation and high sensitivity (multi- copy gene), quantitatively demonstrated by Hachimi (2024) [47], and as a taxonomic standard for establishing phylogenetic relationships [35]. Moreoverm this gene is a tool for the accurate and consistent identification of different *Cryptosporidium* species using a single-primer set [48].

Subsequently, epidemiological subtyping using the gp60 gene may be conducted to explore contamination sources and differentiate between zoonotic and anthroponotic transmission. Our findings are discussed in terms of the potential zoonotic risk of this parasite and as an early warning of its influence in the region.

## Supporting information

https://www.frontiersin.org/articles/10.3389/fvets.2026.2006832/%20full#supplementary-material

## Data Availability Statement

The 18S rDNA sequence generated in this study has been deposited in GenBank under accession number PV442491.1 (https://ncbi.nlm.nih.gov/nuccore/PV442491). Any additional data supporting the findings of this study are available from the co-corresponding authors upon request.

## Ethics statement

All protocols were approved by the Institutional Committee for Animal Care and Use (CICUA) of the University of Chile (Protocol Nº 14-2019, “*Estudio sanitario en castores (Castor canadensis) introducidos en Tierra del Fuego, Chile*”.

## Author Contributions

PL: Investigation, Methodology, Data curation, Visualization, Writing; G. Contreras: Investigation, Methodology, Data curation, Formal analysis; G. Cabrera: Methodology; CA: Resources and management of trapping and sampling; CD: Resources; FF: Conceptualization, Project administration, Resources, Funding acquisition, Supervision, Investigation, Methodology, Writing; CB: Conceptualization, Project administration, Resources, Funding acquisition, Supervision, Investigation, Methodology, Formal analysis, Visualization, Writing.

## Acknowledgements

We extend our gratitude to the administration and the park ranger team of WCS Chile for granting access to Karukinka Park, as well as for their guidance and continuous assistance throughout the research process and the primary collection of biological samples. We also acknowledge the ongoing training, methodological validation, and confirmation of findings, as well as the significant support provided by Alejandra Sandoval and Matilde Larraechea from ConserLab; Claudio Abarca, Sofía Jara and Galia Ramírez-Toloza from the Laboratory of Parasitology, and Fernando Fredes at the Department of Animal Preventive Medicine. Faculty of Veterinary and Animal Sciences, Universidad de Chile. Santiago, Chile.

We are grateful for the invaluable support provided by Séverine Cazaux, Valentina Ferrando, Alejandro Aravena, Javiera Avilés, Ariel Cerda, Tamara Matute, Isaac Núñez, and Dominique Schwend in protein purification, design of synthetic positive controls and PCR/LAMP primers, molecular assays standardization and sample processing. Finally, we acknowledge the conceptual and experimental contributions of Frank Katzer, Hayder Al Mshelesh, Ghulam Musaddaq, Linda Marriott, and Maïwenn Kersaudy-Kerhoas to work on *Cryptosporidium*.

---

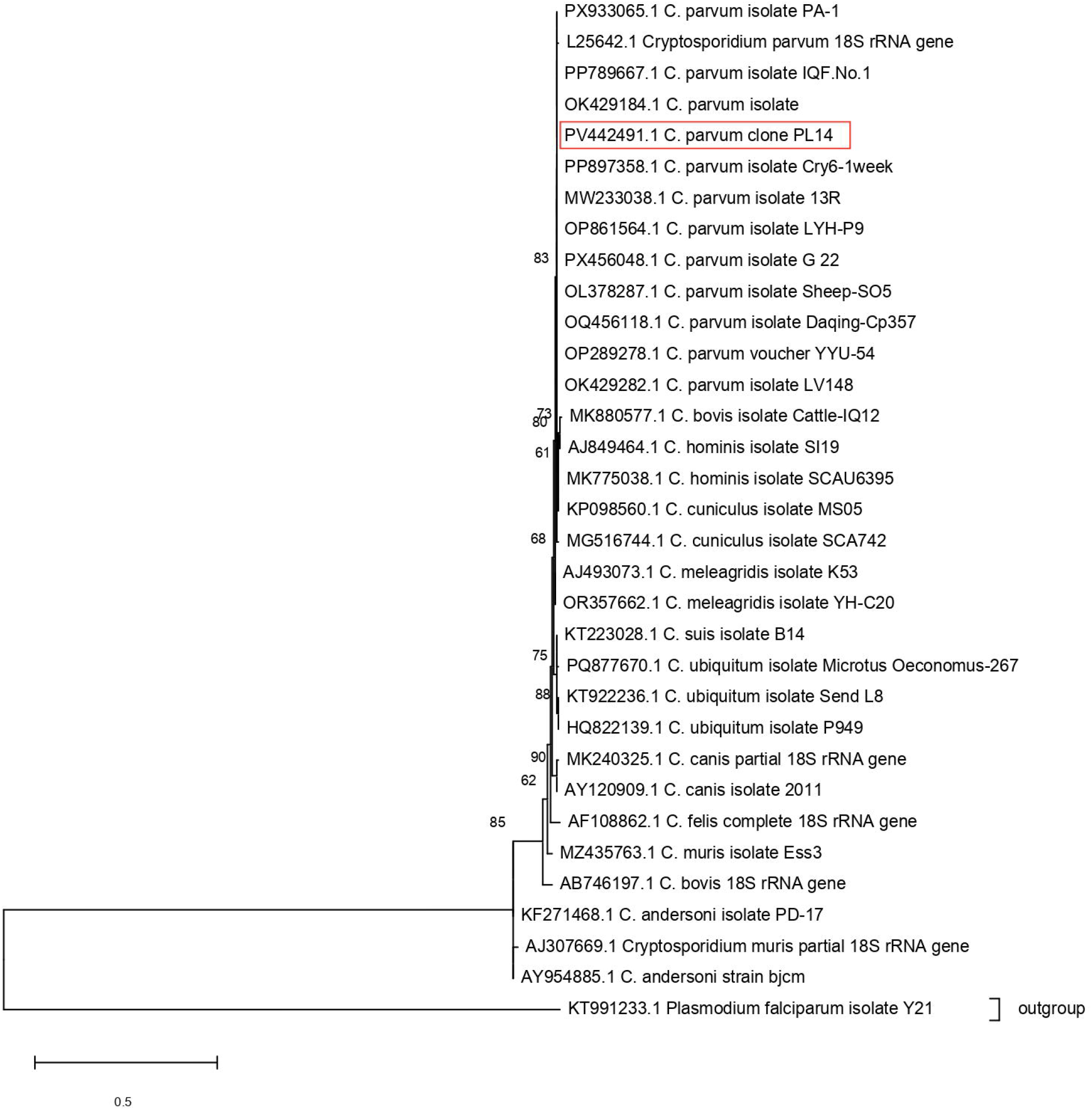

