## Supplementary material for "Detection of zoonotic *Cryptosporidium parvum* in invasive beavers from southern Tierra del Fuego, Chile": https://www.frontiersin.org/articles/10.3389/fvets.2026.2006832/%20full#supplementary-material

### PHYLOGENETIC ANALYSIS

**Supplementary Table 1.** *Cryptosporidium* sequences sourced from GenBank for alignment and interspecies phylogenetic analysis.

| Sequence description | Hosts | Origin | Identity (%) with target |
|---|---|---|---|
| AF513227.2 *Cryptosporidium* sp. strain EGK 3 18S ribosomal RNA gene, partial sequence | Eastern grey kangaroos *Macropus giganteus* | Australia | 100 % |
| KX056094.1 *Cryptosporidium* sp. isolate SSU-rRNAIS13 small subunit ribosomal RNA gene, partial sequence | Humans | India | 100 % |
| KX056097.1 *Cryptosporidium* sp. isolate SSU-rRNAIS16 small subunit ribosomal RNA gene, partial sequence | Humans | India | 100 % |
| L25642.1 *C. parvum* 18S ribosomal RNA (18S rRNA) gene | Newborn calf | Alberta, Canada | 100 % |
| OL378287.1 *Cryptosporidium parvum* isolate Sheep-SO5 small subunit ribosomal RNA gene, partial sequence | Sheeps | Cyprus | 100 % |
| PP897358.1 *Cryptosporidium parvum* isolate Cry6-1week small subunit ribosomal RNA gene, partial sequence | Neonatal calves | Slovakia | 100 % |
| OK429184.1 *Cryptosporidium parvum* isolate LV50 small subunit ribosomal RNA gene, partial sequence | Cattle | Latvia | 100 % |
| OP861564.1 *Cryptosporidium parvum* isolate LYH-P9 small subunit ribosomal RNA gene, partial sequence | Dairy calves | Mongolia, China | 100 % |
| OQ456118.1 *Cryptosporidium parvum* isolate Daqing-Cp357 small subunit ribosomal RNA gene, partial sequence | Dairy calves | Mongolia, China | 100 % |
| MW233038.1 *Cryptosporidium parvum* isolate 13R small subunit ribosomal RNA gene, partial sequence | Wastewater | São Paulo, Brazil | 100 % |
| PP789667.1 *Cryptosporidium parvum* isolate IQF.No.1 small subunit ribosomal RNA gene, partial sequence | Fish and human (fish handlers) | Iraq | 100 % |
| OP289278.1 *Cryptosporidium parvum* voucher YYU-54 small subunit ribosomal RNA gene, partial sequence | Dogs | Turkey | 100 % |
| PX456048.1 *Cryptosporidium parvum* isolate G_22 small subunit ribosomal RNA gene, partial sequence | Dairy calves | Japan | 100 % |

| | | | |
|---|---|---|---|
| PX933065.1 *Cryptosporidium parvum* isolate PA-1 small subunit ribosomal RNA gene, partial sequence | Neonatal calves | Turkey | 100 % |
| OK429282.1 *Cryptosporidium parvum* isolate LV148 small subunit ribosomal RNA gene, partial sequence | Cattle | Latvia | 100 % |
| MK880577.1 *C. bovis* isolate Cattle-IQ12 small subunit ribosomal RNA gene, partial sequence | Cattle and humans | Iraq | 98.46 % |
| AB746197.1 *Cryptosporidium bovis* gene for 18S rRNA, partial sequence, isolate: 21 | Calves | Japan | 93.82 % |
| MK775038.1 *C. hominis* isolate SCAU6395 small subunit ribosomal RNA gene, partial sequence | Equine animals | China | 99.56 % |
| AJ849464.1 *Cryptosporidium hominis* partial 18S rRNA gene, isolate SI19 | Humans | Slovenia | 98.68 % |
| AJ493073.1 *Cryptosporidium meleagridis* 18S rRNA gene, isolate K53 | Humans | Kenya, Malawi, Brazil, UK, Vietnam | 98.45 % |
| OR357662.1 *C. meleagridis* isolate YH-C20 small subunit ribosomal RNA gene, partial sequence | Humans and domestic animals | Heilongjiang Province, China | 98.45 % |
| AJ307669.1 *Cryptosporidium muris* partial 18S rRNA gene | Humans | UK | 90.33 % |
| MZ435763.1 *C. muris* isolate Ess3 small subunit ribosomal RNA gene, partial sequence | Humans, cattle, chickens, sheep and goats | Kenya | 95.10 % |
| AY954885.1 *Cryptosporidium andersoni* strain bjcm 18S ribosomal RNA gene, complete sequence | Dairy cattle | China | 90.33 % |
| KF271468.1 *C. andersoni* isolate PD-17 18S ribosomal RNA gene, partial sequence | Diarrheal human outpatients | China | 90.99 % |
| AF108862.1 *C. felis* 18S ribosomal RNA gene, complete sequence | Humans, cattle, mice, koalas, kangaroos, pigs, cats | South and western Australia, England, USA, Switzerland, Perth | 92.14 % |
| MK240325.1 *Cryptosporidium canis* 18S ribosomal RNA gene, partial sequence | Dogs | São Paulo, Brazil | 90.61 % |
| AY120909.1 *C. canis* isolate 2011 small subunit ribosomal RNA gene, partial sequence | Dogs | USA, Australia, China, Peru, Czech Republic | 96.23 % |
| MG516744.1 *Cryptosporidium cuniculus* isolate SCA742 18S ribosomal RNA gene, partial sequence | Water catchments | Perth, Australia | 98.67 % |
| KP098560.1 *C. cuniculus* isolate MS05 18S small subunit ribosomal RNA gene, partial sequence | Environmental samples | São Paulo, Brazil | 99.34 % |
| KT223028.1 *Cryptosporidium suis* isolate B14 18S ribosomal RNA gene, partial sequence | Pigs | Denmark | 96.48 % |

| | | | |
|---|---|---|---|
| KT922236.1 *Cryptosporidium ubiquitum* isolate Send L8 18S ribosomal RNA gene, partial sequence | Calves and lambs | Central Ethiopia | 97.34 % |
| HQ822139.1 *C. ubiquitum* isolate P949 small subunit ribosomal RNA gene, partial sequence | Surface water, livestock and wildlife samples | North-west England | 97.34 % |
| PQ877670.1 *C. ubiquitum* isolate Microtus Oeconomus-267 small subunit ribosomal RNA gene, partial sequence | Tundra vole | China | 97.56 % |
| KT991233.1 *Plasmodium falciparum* isolate Y21 18S ribosomal RNA gene, partial sequence | Outgroup | China | Not significant similarity |
